# Computation-aided Design of Rod-Shaped Janus Base Nanopieces for Improved Tissue Penetration and Therapeutics Delivery

**DOI:** 10.1101/2024.01.24.577046

**Authors:** Jinhyung Lee, Wuxia Zhang, Danh Nguyen, Libo Zhou, Justin Amengual, Jin Zhai, Trystin Cote, Maxwell Landolina, Elham Ahmadi, Ian Sands, Neha Mishra, Hongchuan Yu, Mu-Ping Nieh, Kepeng Wang, Ying Li, Yupeng Chen

## Abstract

Despite the development of various drug delivery technologies, there remains a significant need for vehicles that can improve targeting and biodistribution in “hard-to-penetrate” tissues. Some solid tumors, for example, are particularly challenging to penetrate due to their dense extracellular matrix (ECM). In this study, we have formulated a new family of rod-shaped delivery vehicles named Janus base nanopieces (Rod JBNps), which are more slender than conventional spherical nanoparticles, such as lipid nanoparticles (LNPs). These JBNp nanorods are formed by bundles of DNA-inspired Janus base nanotubes (JBNts) with intercalated delivery cargoes. To develop this novel family of delivery vehicles, we employed a computation-aided design (CAD) methodology that includes molecular dynamics and response surface methodology. This approach precisely and efficiently guides experimental designs. Using an ovarian cancer model, we demonstrated that JBNps markedly improve penetration into the dense ECM of solid tumors, leading to better treatment outcomes compared to FDA-approved spherical LNP delivery. This study not only successfully developed a rod-shaped delivery vehicle for improved tissue penetration but also established a CAD methodology to effectively guide material design.

## Introduction

A key limitation in current treatment strategies is the lack of specificity in delivery systems targeting tissue and the low efficacy of therapeutics due to the dense and complex extracellular matrix (ECM) of tissue [2, 3]. This is particularly evident in certain solid tumors, like ovarian cancer, where the ECM not only creates a protective environment but also acts as a formidable barrier to efficient drug penetration [4, 5]. Therefore, overcoming this barrier is essential for effective therapy. Currently, lipid nanoparticles (LNPs) are the most commonly used delivery vehicles; however, their size range of 50-500 nm limits their penetration capability in tissues characterized by a dense ECM. Consequently, the development of innovative biomaterials that can enhance the potency of chemotherapy drugs is a critical step toward advancing cancer treatment. In this context, the development of novel nanomaterials marks a pivotal advancement in delivery methods [6, 7].

We introduce a new family of nanomaterial: Janus base nanomaterials (JBNs), self-assembled from a library of small molecules. These comprise DNA-mimicking bases (guanine and cytosine) and an amino acid side chain. The chemistry, structure, and formation of JBNs are distinct from any conventional nanomaterials such as lipids, polymers, carbon nanotubes and DNA origami. JBNs can self-assemble into noncovalent nanotubes (called JBNt) based on hydrogen bonds and *π*-*π* interactions. Previously, we showed that a delivery platform based on JBNts could effectively deliver small RNAs into the cell via enhanced endosomal escape through the “proton-sponge” effect with excellent biocompatibility [32]. In this study, we have pioneered the use of DNA-inspired Janus bases as a delivery platform, employing computation-aided design (CAD) for its development.

The traditional methods for formulating nanoparticles for drug delivery have often relied on a resource-intensive and time-consuming process, commonly referred to as the “Edisonian” approach. This method is predominantly characterized by trial-and-error, lacking a systematic and theoretical framework. Recent advancements in high-fidelity Computer-Aided Design (CAD) have become crucial in providing in-depth insights into the properties and behavior of nanomaterials, significantly improving their effectiveness and efficiency in drug delivery systems. This new methodology comprises three key steps: (1) the employment of molecular dynamics (MD) simulations to analyze the surface chemistry of JBNts, (2) the investigation of the potential of mean force (PMF) in relation to the loading of cargoes into JBNt, and (3) the use of response surface methodology (RSM) to optimize the encapsulation efficiency of these cargoes. This approach not only reduces the reliance on trial-and-error but also enhances the precision and predictability of designing novel drug delivery systems.

In this work, we present a streamlined method using CAD to simulate the self-assembly of Janus base Nanotubes (JBNt) and the incorporation of cargoes within them. This method was not only theoretically conceptualized but also experimentally validated, along with an evaluation of the cargo delivery capabilities. Subsequently, we successfully transformed the non-rod-shaped JBNt nanoparticles (Non-rod JBNp) into a rod-shaped JBNt nanoparticles (Rod JBNp). Compared to conventional spherical nanoparticles, such as LNPs, Rod JBNp exhibits superior ability to penetrate the dense ECM of solid ovarian tumors, highlighting its promising effectiveness in these types of applications.

## Results

### Simulation and Experimental Validation of JBNt drug loading

JBNts are self-assembled from a library of nanotube monomers which consist of two components: a base component mimicking DNA bases and an amino acid side chain. Cargoes (DOX in this case) are encapsulated within the JBNt through intercalation between JBNt monomers (**Fig 1A**). To investigate JBNt self-assembly, two key geometrical parameters, including the vertical distance and the relative torsion angle between two G^C Janus base monomers, have to be controlled (**Fig 2A, i**). **Fig 2A, ii** shows 2D-potential energy surfaces of relative stacking energies between two G^C Janus base monomers as a function of the vertical distance and the in-plane relative torsion angle. **Fig 2A, iii** displays the representative configurations of the stacked G^C Janus base monomers, along with their locally optimal vertical distances and in-plane relative torsion angles. The relative stacking energy of G^C Janus base monomers with respect to the vertical distance and relative torsion angle reveals one optimal stacking minimum at about 4 Å and about 30°, respectively.

**Fig. 1:**
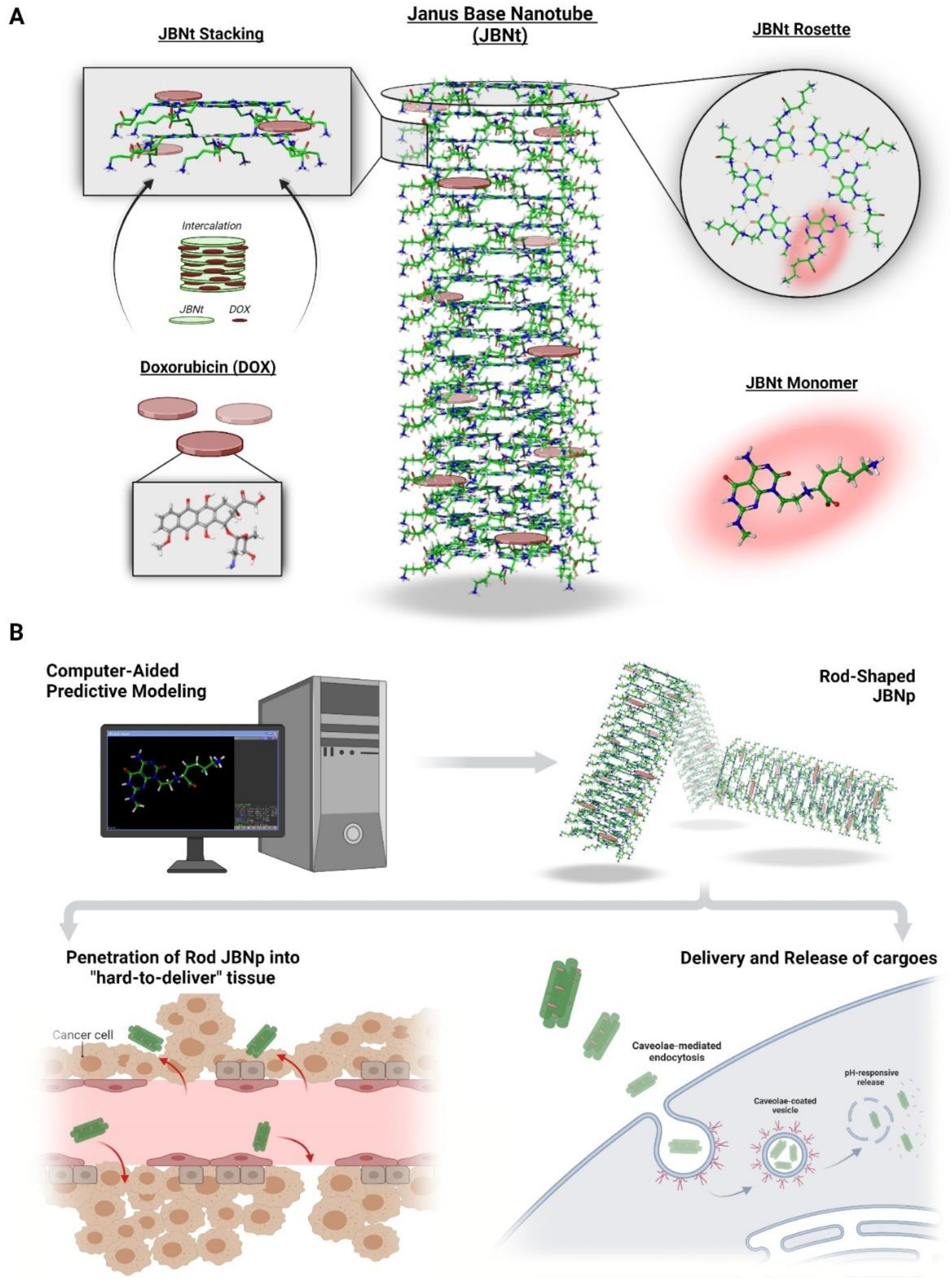
Schematic illustration of the Janus Base Nanopieces (JBNps) for drug delivery. (A) Schematic illustration of self-assembly of Janus base Nanotube (JBNt) and its components. (B) The scheme shows that utilizing the computer-aided design (CAD) predictive modeling to assemble JBNt-cargoes and engineering to make rod shaped nanoparticle, called Rod JBNp. As a proof-of-concept, Rod JBNp delivers and releases cargoes and penetrate “hard-to-penetrate” tissue such as a solid tumor.

**Fig. 2:**
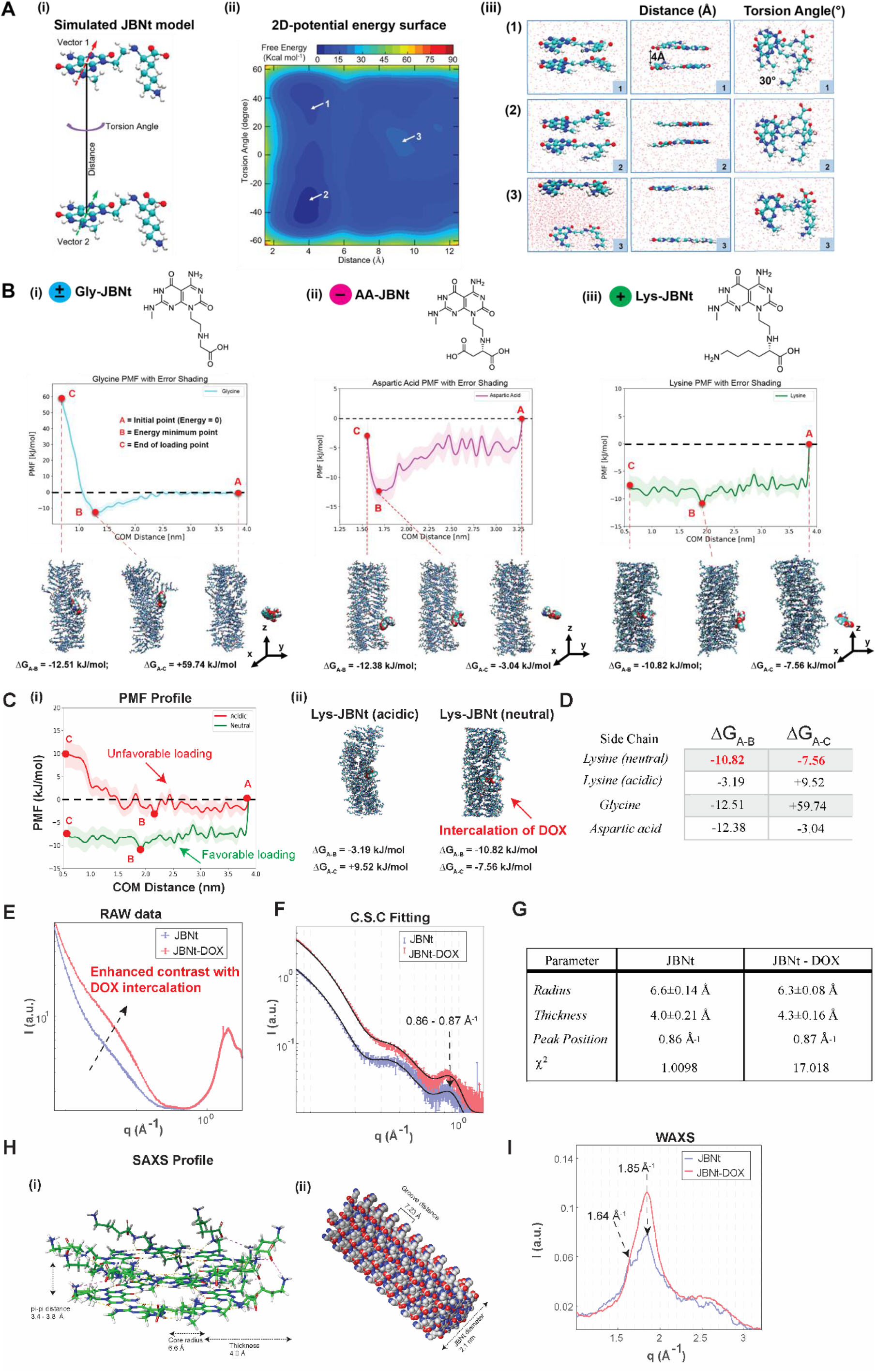
Simulation and Experimental Validation of JBNt drug loading. (A, i) Metadynamics simulation of stacking two G^C Janus base monomers considering the distance and torsional angle between two monomers. (A, ii) 2D potential energy surface of the stacking of two monomers. (A, iii) Representative snapshots of stacking of two monomers. (B, i) Potential of mean force (PMF) of DOX loading onto Glycine-JBNt (Gly-JBNt). (B,ii) PMF of DOX loading onto Aspartic acid-JBNt (AA-JBNt). (B,iii) PMF of DOX loading onto Lysine-JBNt (Lys-JBNt). (C,i) PMF of DOX loading onto Lysine-JBNt in neutral pH and acidic pH. (C, ii) Representative snapshots of intercalation of DOX at different pH values. (D) Table of PMF profiles of DOX loading onto different JBNts. (E) Enhanced intensity due to DOX intercalation at identical Lys-JBNt concentrations. (F) SAXS data and their best fits using the core-shell cylinder and gaussian peak combined model. (G) Tabulated best fitting parameters to the SAXS data. (H) Detailed local structure of Lys-JBNt (I) Two high-q peaks corresponding to the well-defined interlayer spacings of Lys-JBNt monomers.

We then modeled the intercalation of DOX into JBNts. We selected three JBNt candidates from our library: Gly-JBNt, AA-JBNt, Lys-JBNt based on the surface charge of the JBNt, showing three different surface charges of side chain: glycine (neutral), aspartic acid (negative), and lysine (positive) charge, respectively. We then calculated the potential of mean force (PMF) of each JBNt involved in the intercalation of DOX. **Fig 2B i, ii, iii** illustrates the PMF profiles of each JBNt involved in the intercalation of DOX. Among the JBNt candidates, Lys-JBNt displays the lowest Δ*G*_*A*−*B*_ = −10.82 kJ/mol, and Δ*G*_*A*−*C*_ = −7.56 kJ/mol, indicating the relatively lower energy barriers for both the absorption of DOX into the side chain of JBNt and for the intercalation of DOX within JBNt compared to Gly-JBNt and AA-JBNt. As a result, Lys-JBNt was selected for DOX loading.

We further investigated the binding capacity of the DOX molecule on the JBNt under various pH conditions including neutral (high pH) and acidic (low pH) conditions. **Fig 2C, i** illustrates the PMF profiles of Lys-JBNt and DOX molecule in the water phase at different pH levels. The calculated free energy of the system gradually decreases as DOX approaches the center of the JBNt, indicating a favorable DOX intercalation process in the neutral environment. Conversely, DOX encounters challenges when entering the center of JBNts under low-pH conditions. We then calculated the DOX loading energies under different pH levels from the PMF profiles. By subtracting the final loading energy from the initial loading energy, we determined binding energies of −7.56 kJ/mol and 9.52 kJ/mol for DOX to Lys-JBNts under neutral and acidic conditions, respectively (**Fig 2C, ii**). These binding energy values confirm the more favorable loading of DOX onto Lys-JBNts under neutral conditions in contrast to low-pH conditions. All the energies associated with the loading of DOX onto different JBNts are summarized in **Fig 2D**.

**Fig 2E** indicates intensity enhancement after the loading of DOX. The reduced SAXS data (after transmission and background correction) were analyzed and best fitted via SasView software using a model that combines a core-shell cylinder (CSC) with a single gaussian peak (**Fig 2F**). The best fits using this combined model agree well with the SAXS data. The CSC model suggests that the JBNt forms a hollow rod structure, consistent with the proposed structure and the shell of the cylinder represents the Lys-JBNt. From the fitting results (**Fig 2G**), the Lys-JBNt diameter is found to be conserved ∼ 2.1±0.24 nm, suggesting that the radial dimension of the Lys-JBNt undergoes minimal variation after DOX loading. It is noteworthy that electron densities of the core (ρ_c_), shell (ρ_s_), and water (ρ_w_) are in the order of ρ_s_ > ρ_c_ > ρ_w_ consistent with the calculated electron densities. The best fitted scattering contrast (the difference in electron density) between core and water decreases while that between shell and water increases after DOX loading. It indicates that DOX (with a higher electron density) is associated with the Lys-JBNt shell resulting in a higher electron density. A small portion of DOX can also localize near the core, thus increasing ρ_c_ and reducing the difference between ρ_c_ and ρ_w_. Intercalation of DOX enhances the total contrast between the CSC and solvent leading to the intensity enhancement seen at equivalent Lys-JBNt concentration (**Fig 2E**).

The best-fitted Gaussian peak position locates at 0.87Å^−1^, corresponds to a regular repeat spacing of 7.2 Å. The spacing can be rationalized by detailed local structure of the Lys-JBNt (**Fig 2H**). **Fig 2I** shows the wide-angle X-scattering (WAXS) data of the two samples (in the presence and absence of DOX). Both show two peaks at 1.64 and 1.85Å^−1^, corresponding to the spacings of 3.83Å and 3.4Å, respectively, which are in good agreement with the simulation results of the Lys-JBNt groove distance: the vertical length between parallel lysine groups after a full rotation of Lys-JBNt rosettes. Sum of the two spacings yields a length of 7.23Å, consistent with the corresponding length of the Gaussian peak (7.2 Å), implying that 3.83 Å and 3.4 Å are most likely alternating vertical spacing between the Lys-JBNt rosette. Therefore, for the first time, we have successfully conducted a simulation of the JBNt to determine the optimal side chain and condition to intercalate small molecules, confirming our findings with exceptional precision through SAXS analysis.

### RSM-based computed method to optimize the JBNps

To validate the simulation, the UV-visible (Vis) spectra demonstrated molecular-level incorporation between JBNts and DOX. When assembled with DOX, the 280-nm peak of JBNt-DOX decreased due to the intercalation between JBNt units and DOX. At pH 8.3, 38 °C and 72h, a significant decrease of absorbance at 280 nm was observed compared to controls, demonstrating that DOX was able to load into JBNts successfully under explored simulated conditions (**Fig 3A, 3B, 3C**). Then, we assessed the encapsulation efficiencies (EE) across varying pH through nuclear magnetic resonance (NMR) (**Fig S6A**). **Fig 3F** shows the NMR-based calculated EE with three factors (pH, temperature, and time). The results correspond to the UV-Vis trends. To enhance the optimization of JBNt-DOX, we utilized a computed approach based on RSM. **Fig 3D (i, ii, iii)** presents the 2D RSM contour plots, demonstrating the incorporation of DOX into JBNt under varying conditions such as temperature, pH, and time. Additionally, the 3D RSM surface plots are depicted in **Fig 3E (i, ii, iii)**. Collectively, Figure 3G illustrates a 3D cube computation, incorporating the three aforementioned factors in relation to the EE. Finally, based on the RSM, we have maximized the EE to 93 % of DOX into JBNt showing significant decrease of UV-Vis absorbance (**Fig S6**) and imaged the JBNt-DOX that turned to saturated red color in PBS (**Fig 3H**). As an initial demonstration of JBNt ability to incorporate cargoes into JBNt, we also effectively loaded resveratrol, a promising antioxidant and anticancer drug, into JBNt **(Fig S8)**.

**Fig. 3:**
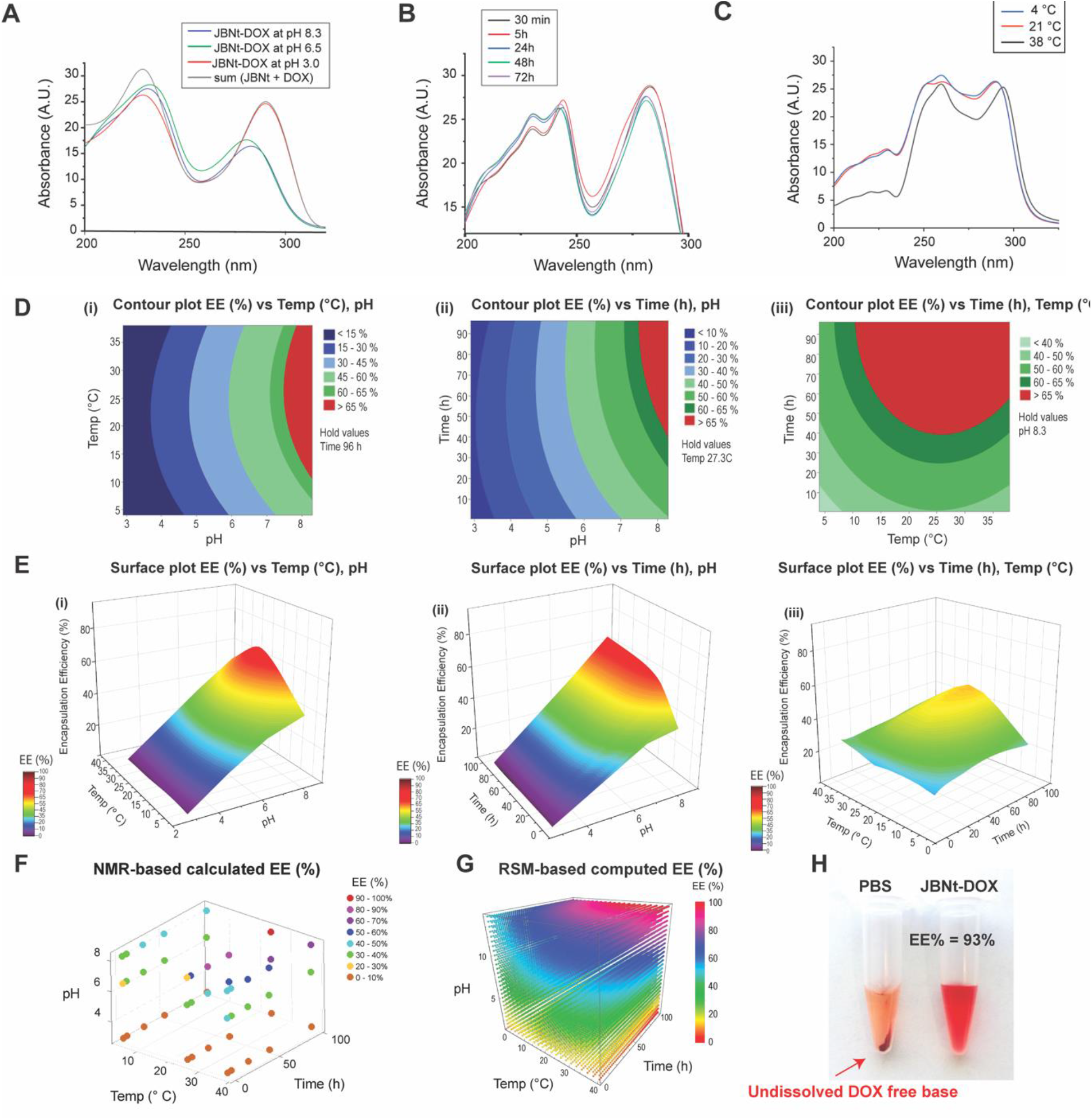
Optimization of JBNp formulation based on RSM-computed method. (A) UV-Vis analysis of pH-dependent assembly. (B) UV-Vis analysis of time-dependent assembly. (C) UV-Vis analysis of temperature-dependent assembly. Response surface methodology of DOX loading to JBNt. (D) 2D RSM contour plots for the EE(%) with different conditions; (i) temp, pH. (ii) time, pH. and (iii) time, temp. (B) 3D RSM surface plots for the EE(%) with different conditions; (i) temp, pH. (ii) time, pH. and (iii) time, temp. (F) NMR-based calculated encapsulation efficiency (EE %). (G) 3D cube figure of RSM-based computed EE %. (H) Photograph of solution of DOX (left) and JBNt intercalated DOX (right) in the ambient light.

### Fabrication and characterization of rod shaped JBNps

Guided by MD simulations, we can further process JBNt with DOX molecules into delivery vehicles. JBNt alone presents nanotubular morphology (length of ∼ 300nm) **(Fig 4A, i** and **Fig S10A)**, forming non-rod-shaped nanoparticle after incorporating with cargoes **(Fig 4A, ii**). Interestingly, a simple sonication process can break these bundles into smaller individual rod-like vehicles (named Rod JBNp, **Fig 4A, iii**). Although the whole JBNp architecture is formed by noncovalent interactions of their small molecule units and cargoes, the JBNps are stable entities. As shown in **Fig 4B** and **Fig S11B,** we altered the sonication power to process Non-rod JBNp to Rod JBNp. Their aspect ratio (AR), the length (k) divided by the width (*k*_*p*_) increases as sonication power (%) increases. The Rod JBNp’s AR were 5.3, which has length of 126.3 ± 13.9 nm, and its width of 26.7 ± 2.5 nm. In accordance, the zeta potential measurements of JBNp demonstrated the shift of their surface charge by sonication power (**Fig 4C**). Then, we fixed the sonication power to 100% and alter the sonication time. As shown in **Fig S10C** and **Fig S10D**, 150 seconds of sonication time formed the Rod JBNp with surface charge of ∼ 7 mV. UV-Vis spectra demonstrated there is a dissociation between JBNps and DOX when decreasing the pH from 7.4 to 5.2 **(Fig 4D**). Moreover, as shown in **Fig 4E**, Rod JBNp dissociated and released the DOX shown in agarose gel electrophoresis. The apparent pKa of Rod JBNp was determined as ∼7.65 by the 2-(p-toluidino)-6-naphthalene sulfonic acid (TNS) binding assay, which indicates the optimal range for tumor delivery and escape from endosome via the “proton sponge” effect **(Fig 4F**). In addition, we evaluated the cytotoxicity of the JBNps. Relative cell viabilities of fibroblast cells after incubating with delivery materials by increasing concentrations. Compared with various commonly used delivery materials: Lipo, SWCNTs, PEI, PLL, the JBNps showed significantly better cell viability (**Fig 4G**).

**Fig. 4:**
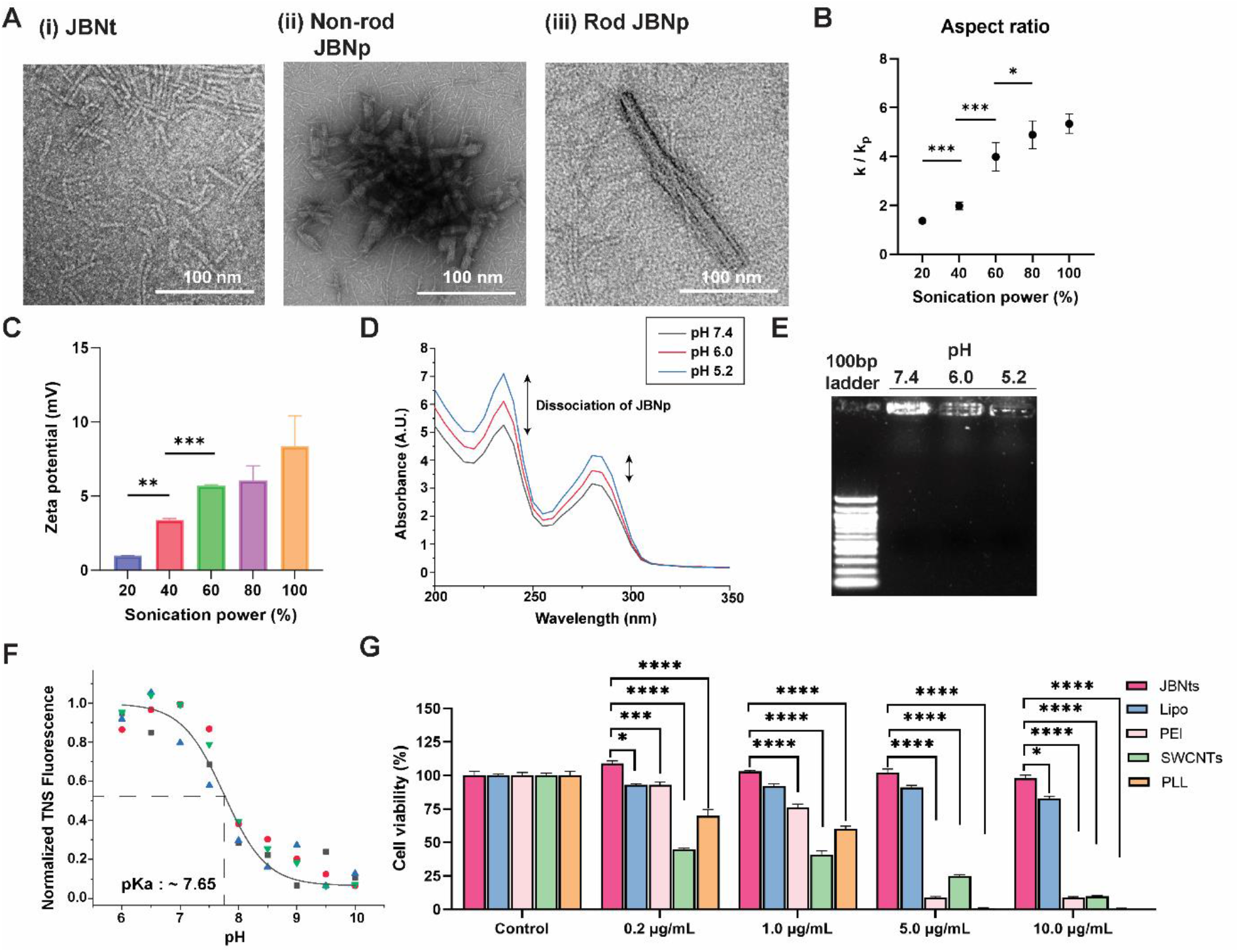
Fabrication and characterization of rod shaped JBNps. (A) TEM images of JBNts, Non-rod JBNp, and Rod JBNp. (B) Aspect ratio analysis. (C) Zeta potential analysis varying sonication power. n=3. (D) UV-Vis analysis. (E) Agarose gel electrophoresis assay varying pH. (F) Apparent pKa assay of Rod JBNp. n=3. (G) *In vitro* cytotoxicity assay: Lipofectamine2000 (Lipo), single-walled carbon nanotubes (SWCNTs), polyethyleneimine (PEI), and poly-L-lysine (PLL). The data were expressed as the percentage of surviving cells, and the values are mean ± SEM (n≥ 10). *P < 0.05, ** P < 0.01, and *** P < 0.001 compared to untreated control.

### *In vitro* delivery and functional assay of rod shaped JBNps in cancer cells and spheroids

Based on their excellent drug loading capacity, Rod JBNp exhibit a great potential to be delivery vehicles for drugs into cells. Thus, we demonstrated the intracellular delivery ability of the Rod JBNp via confocal laser scanning microscopy (CLSM) images. Rod JBNp was able to deliver DOX to cell in a time-dependent fashion, showing red signal (DOX) in cytoplasm (**Fig S12**). As depicted in **Fig 5A, i, Fig S13**, ovarian cancer (SKOV3) cells and breast cancer (MCF7) cells presented higher DOX (red) fluorescent intensity than the non-rod JBNp–DOX and DOX-freebase groups. CLSM z-stack images showed that DOX (red) successfully delivered to SKOV-3 cells (**Fig S13**). Not only can we deliver DOX, but we can also deliver messenger RNA (mRNA) cargoes to human chondrocytes (C28/I2 cells), as shown in **Fig. 5A, ii**. This demonstrates the versatility of Rod JBNp in delivering various kinds of cargoes. As presented in **Fig 5B, Fig S13B**, the mean DOX intensity of the Rod JBNp group is significantly higher than the other groups, which indicated that DOX can be delivered into the nucleus more effectively by JBNp as compared to the controls. In addition, the mean fluorescence intensity of DAPI for each group was also statistically analyzed. As indicated in **Fig 5C, Fig S13C**, the fluorescent intensity of DAPI in the Rod JBNps–DOX group was significantly weaker than that controls, which may be caused by the failure of DAPI binding with damaged DNA.

**Fig. 5:**
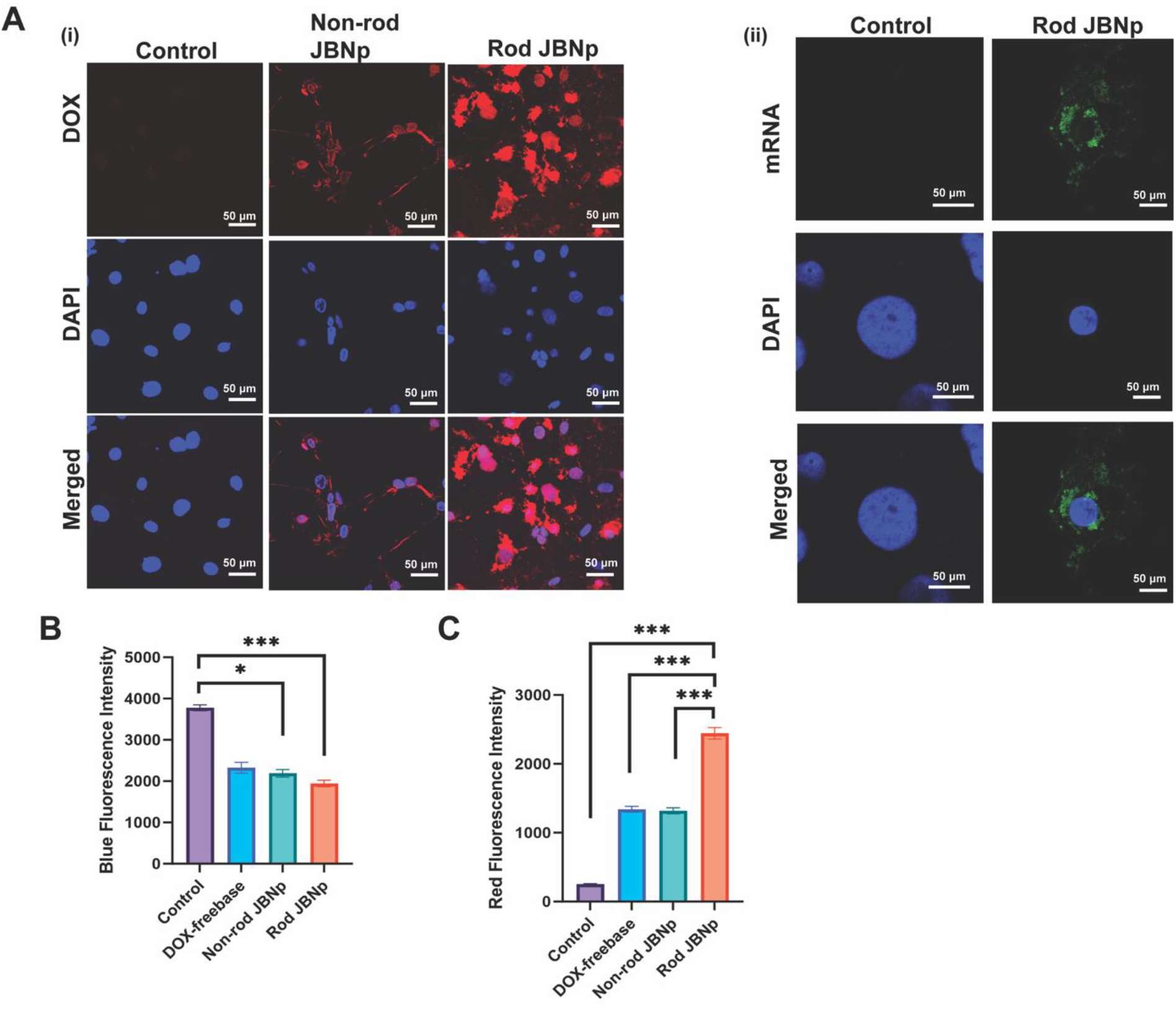
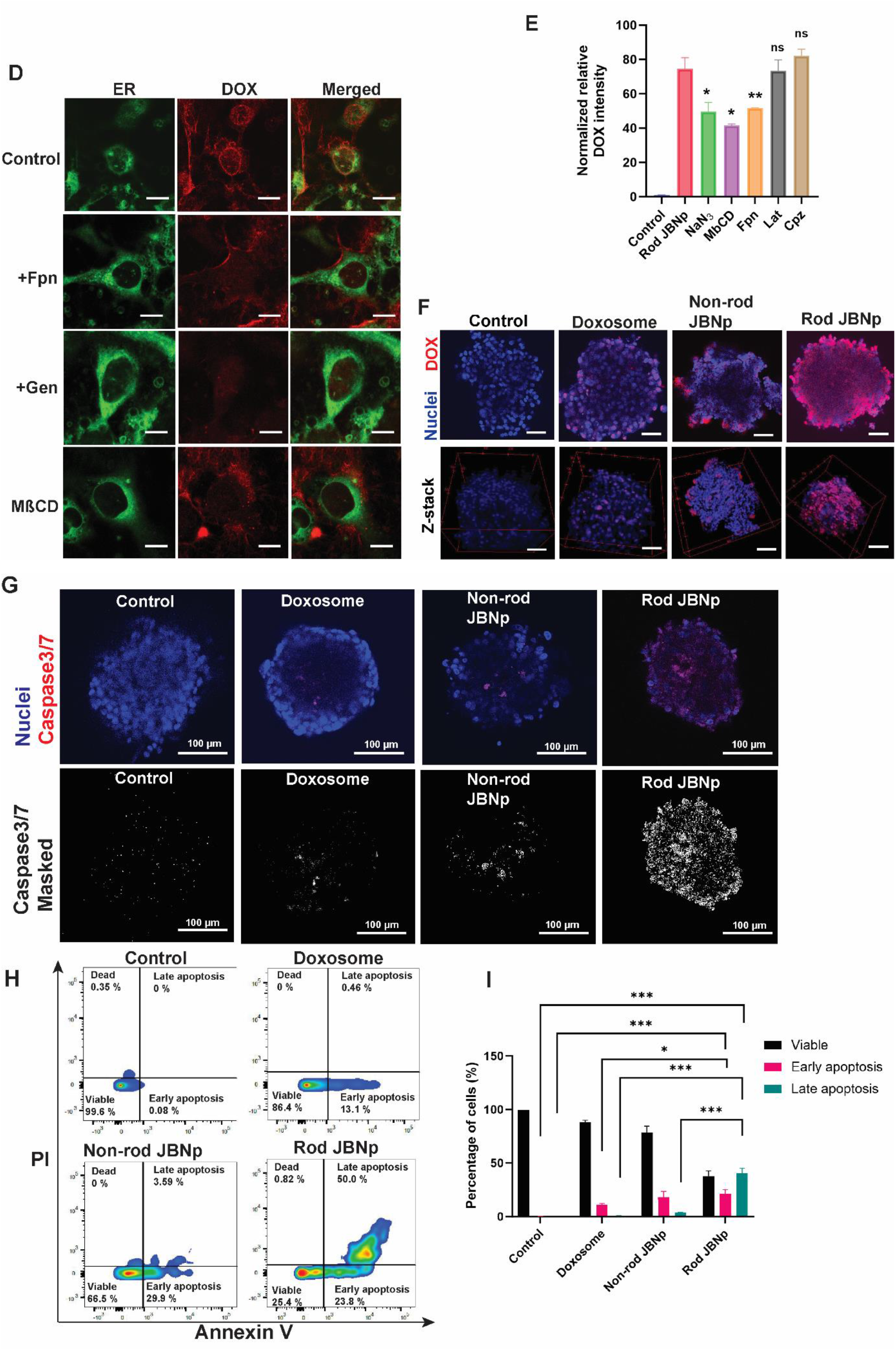
Rod shaped JBNps showed efficient delivery to cells and cancer spheroid. (A, i) CLSM images of DOX (red) delivered by the DOX-freebase, Non-Rod JBNp, and Rod JBNp. (A, ii) CLSM images of Cas9-eGFP mRNA (green) delivered by the Rod JBNp. (B) Statistical analysis of red fluorescence intensity of confocal images**. (**C) Statistical analysis of mean blue fluorescence intensity. (D) CLSM images of Rod JBNp pretreated with caveolae-mediator inhibitors; Endoplasmic reticulum (green); DOX (red). (E) Quantitative analysis of Rod JBNp uptake. (F) CLSM images of SKOV3 spheroids delivered by the Rod JBNp; cell nuclei stained with DAPI (blue); DOX (red). (G) CLSM images of SKOV spheroids treated with indicated group. Stained with Hoechst (blue) and Caspase 3/7 (red) reagents.; Apoptotic cells are identified by a white mask. (H) Apoptosis assay of SKOV-3 spheroids after treatments for 48h using Apoptosis Kit with Annexin V FITC and PI. (I) Percentage of viable, early apoptotic, and late apoptotic cells. n=3. Data are presented as the means ± SEM of triplicate experiments. *P<0.05, **P<0.01, and ***P<0.001. Data are presented as the means ± SEM. N=5.

Moreover, because the JBNps have different chemical compositions compared to conventional delivery materials, we also explored the Rod JBNp cellular uptake mechanism. We pretreated the SKOV-3 cells with several endocytic inhibitors, including ATP inhibitor (NaN_3_), caveolae inhibitor (Genistein [Gen], methyl-*β*-cyclodextrin [*MβCD*], a fillipin [Fpn]), clathrin inhibitor (chlorpromazine [Cpz]), and a macropinocytosis inhibitor (latrunculin A [Lat]). The CLSM image of qualitative results in **Fig 5D** demonstrated significant inhibition of Rod JBNp uptake and delivery to the endoplasmic reticulum (ER) using caveolae-mediated inhibitors (Fpn, Gen, and *MβCD*). Similarly, the quantitative results presented in **Fig 5E** demonstrated the same significant inhibition of uptake when pre-treating with caveolae-mediated inhibitors. Thus, the results suggest that the uptake mechanism of Rod JBNps–DOX relies on the energy dependent caveolae-mediated transcytosis. Moreover, the *in vitro* tumor transcytosis of Rod JBNps–DOX was further investigated by their delivery to the ovarian cancer spheroid. As depicted in **Fig 5F** and **Fig S16**, Rod JBNp–DOX was successfully delivered to cells inside the SKOV3 spheroid compared to peripheral delivery of Doxosome and Non-rod JBNp.

Further assessment of functional assay with Rod JBNp on SKOV-3 cancer cells and spheroid was performed using Caspase 3/7 staining. Caspase 3/7 activity did significantly increase after 24 h of treatment with Rod JBNp when compared to untreated control in SKOV-3 cells (**Fig S17**). Then, annexin V-FITC/Propidium iodide (PI) double staining analysis revealed in **Fig S18,** following treatment with Rod JBNp–DOX for 24h, the percentage of early apoptotic SKOV-3 cells increased to 73.6%. We then demonstrate that the Rod JBNp shows a significantly higher rate of apoptosis by time-dependent manner (**Fig S19**). To further test the functional delivery of DOX, we tested the formation of cancer spheroid upon Rod JBNp. Microscope images and quantitative spheroid size analysis revealed that Rod JBNp–DOX mediates the initial formation of ovarian carcinoma spheroids better than controls (**Fig 5D**).

Finally, we evaluated the apoptosis phenotype in the cancer SKOV-3 spheroid. As shown in **Fig 5G**, the treatment of the spheroid with Rod JBNp–DOX led to a significant increase in the number of apoptotic cells (Caspase 3/7 dye: red) and even indicated an apoptosis signal in the deep center of the spheroid compared to the controls. To quantify the apoptotic signal in the spheroid, high throughput flow cytometry analysis was performed. (**Fig 5H**). The results revealed that the Rod JBNp–DOX achieved an apoptotic efficiency of 61.8%, demonstrating a higher apoptotic effect on the tumor spheroids when compared to the DOX-free base, Doxosome, and Non-rod JBNp groups which demonstrated efficiencies of 9.76%, 11.4% and 21.2%, respectively (**Fig 5I**). This may be due to the rod-shaped morphology of the Rod JBNp, allowing the nanoparticles to penetrate deep into the spheroid and deliver the DOX.

### *In vivo* biodistribution, and Tumor Penetration

Given that rod-shaped JBNps is superior in transcytosis and penetration across densely packed cancer spheroids, we next seek to test its tumor-penetration potential and biodistribution *in vivo*, via tumor xenograft mice. It is known that nanoparticles with an aspect ratio of 5-10 can accumulate in tumor sites due to the penetration of thick ECM. Thus, we investigated the *in vivo* biodistribution assay of Rod JBNps after intravenous injection to determine their tumor specificity and delivery efficiency. The biodistribution of Rod JBNps were evaluated in nude mice bearing a subcutaneous SKOV-3 xenograft via intravenous injection. *Ex vivo* fluorescence intensity images were obtained for tumors and major organs to explore the DOX distribution. As depicted in **Fig 6A**, Rod JBNps showed enriched distribution to the tumor compared to other major organs. The intensities of fluorescence signals in tumors were stronger than those in other major organs. These results were further confirmed by quantitative analysis (**Fig 6B**). Delivery of Rod JBNp has demonstrated significantly higher accumulation of DOX compared to Doxosome and Non-Rod JBNp. Further, to validate the delivery of DOX, the tumor tissues were collected and sectioned for fluorescence examination at 72h (**Fig 6C**). The results revealed that Rod JBNp successfully delivered DOX more deeply into the tumor compared to Doxosome and Non-rod JBNp. Additionally, the distribution of DOX in tumors was analyzed via anti-CD31 immunohistochemistry. Confocal imaging of the stained sections revealed that DOX delivered by Rod JBNp carriers was mainly distributed to the CD31-stained blood vessels in the tumor region (**Fig 6D**). By the passive targeting of tumors via the penetration effect due to rod shaped morphology, JBNp holds the potential to increase the efficacy of drugs while reducing the nonspecific distribution of drug during the *in vivo* delivery process.

**Fig. 6:**
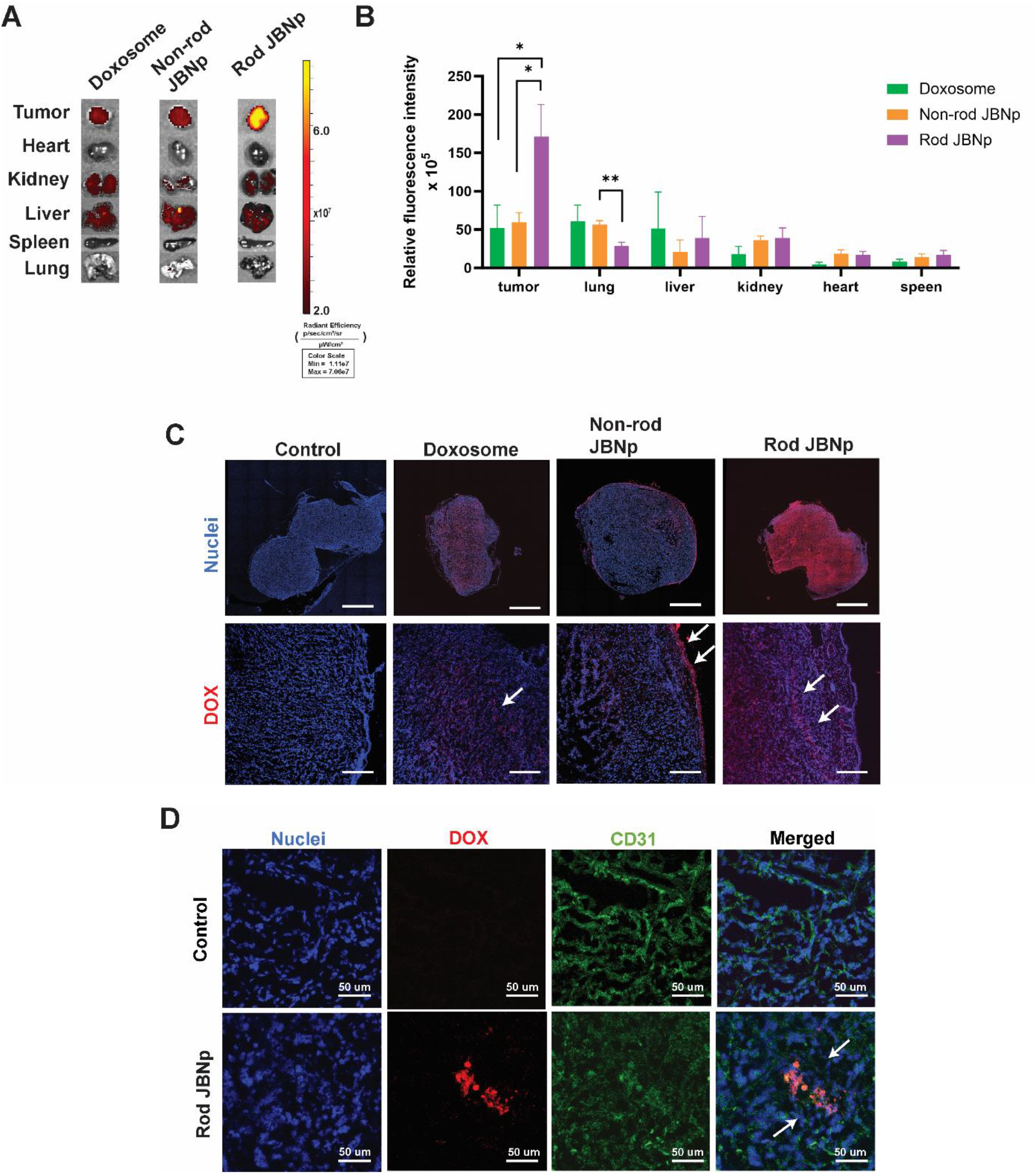
Rod shaped JBNps showed tumor-specific biodistribution and higher delivery efficiency than spherical Doxosome. (A) *ex vivo* biodistribution of major organs and tumors after intravenous injection to SKOV3 tumor xenograft NU/J mice for 72h. (B) *Ex vivo* quantification of relative fluorescence intensity in major organs and tumors. (C) Tumor section and stained with DAPI, Blue: nuclei staining, Red: DOX. (D) Rod JBNp localization in tumor tissue in relation to the tumor blood vessels detected by the CD31 biomarker. Red: DOX; Blue: DAPI staining Green: CD31. The animal was sacrificed at 72h post injection. Error bars, mean ± SEM (n=5).

### Antitumoral effect of rod shaped JBNps to treat SKOV-3 tumor-bearing mice

Based on the encouraging results found in our biodistribution study, we further investigated the *in vivo* antitumor efficacy of Rod JBNp. When the tumor volume in the SKOV-3 tumor-bearing nude mice model reached approximately 200 mm^3^, nude mice harboring these xenographs were randomly divided into four groups: saline, Doxosome, and Rod JBNp. Mice were then intravenously injected every 3 days for 21days. Tumor volume was recorded every three days. After treatments, all mice were euthanized, and collected tumors were weighed and used for the quantitative analysis of the inhibition of tumor growth (**Fig 7A**). As displayed in **Fig 7B**, tumor growth of the saline and Doxosome group were rapid, whereas tumor growth of the Rod JBNp group was significantly inhibited during 21 days of treatment, thereby indicating that Rod JBNp was effective for cancer treatment. The tumors were excised, photographed (**Fig 7C**), and weighted. Compared with saline, Rod JBNp treated mice showed a 59.2 ± 10.2% decreases in tumor weight (**Fig 7D**). Moreover, the H&E staining results of tumor sections showed that there was an increase in necrosis and apoptosis in tumor cells of the Rod JBNp group, compared with that observed in other groups. To further investigate the role of apoptosis, tumor sections were stained for the expression of cleaved caspase-3 and TUNEL in order to detect early and late apoptosis, respectively. (**Fig 7E**). Consistent with the reduction in tumor growth, there was also a significant increase in the expression of the early apoptosis marker, cleaved caspase-3, as compared to treatment with Doxosome (**Fig 7F**). Tumors treated with Rod JBNp also showed a significantly higher intensity of the TUNEL staining, thereby indicating significantly more late apoptotic cells (as indicated in green fluorescence) in tumors than those in other groups (**Fig 7G**). To exclude the inflammation effect in apoptosis, we have also confirmed that the proinflammatory responses in tumor microenvironment are not playing a role in therapy response (**Fig S21**). In conclusion, Rod JBNp was able to penetrate deep into the tumor, delivered DOX and induced significantly higher antitumor activity than Doxosome.

**Fig. 7:**
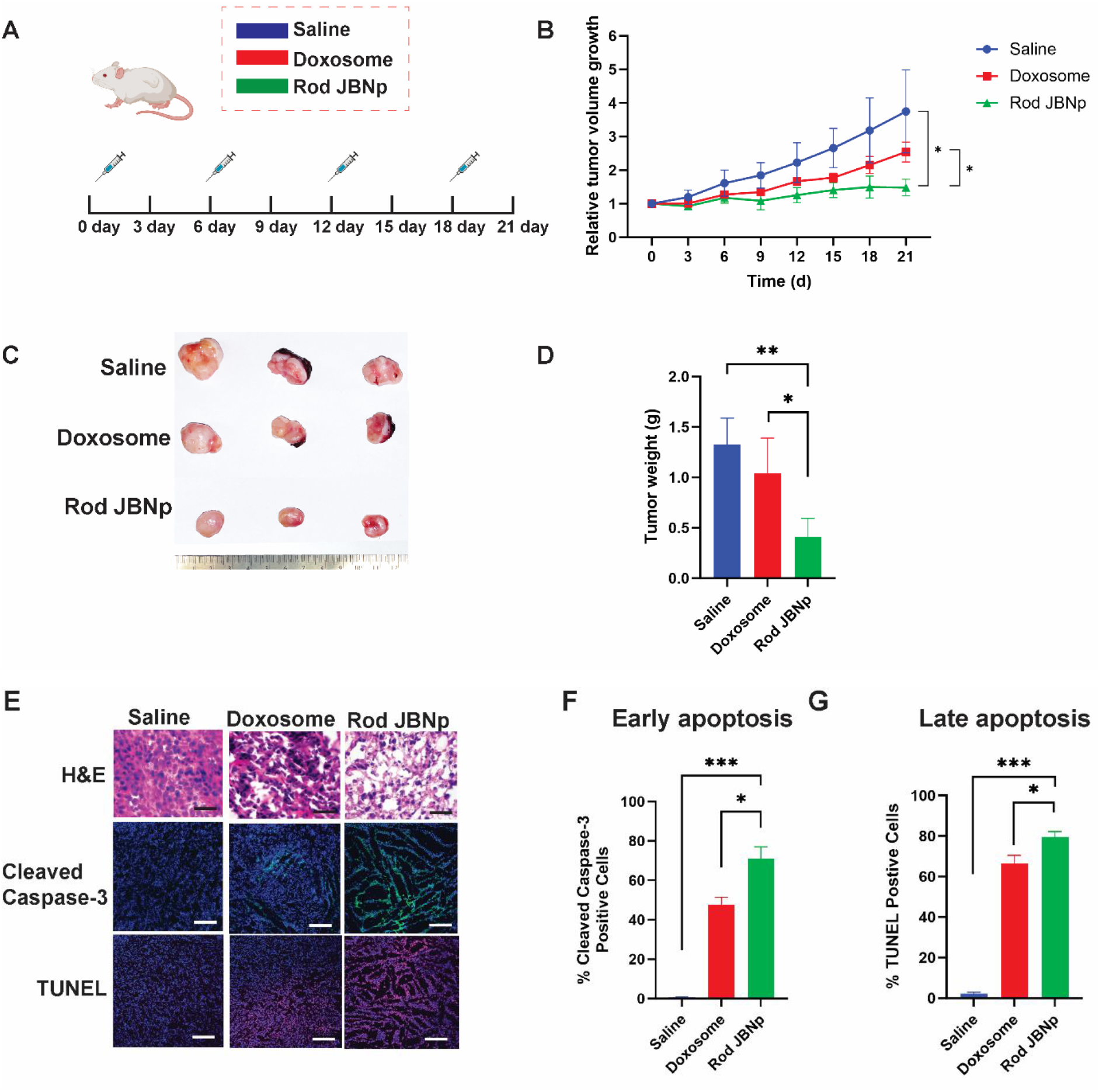
Rod shaped JBNps induced significantly high antitumor activity. (A) Schematic illustration of the treatment plan. (B) Relative tumor-growth curves during the treatment measured by a caliber (n=3 or n=5). (C) Photograph of excised tumors from SKOV-3 tumor-bearing mice after varous treatments. (D) Weight of excised tumor (n=3). (E) Histopathological analysis of the tumors by haematoxylin and eosin (H&E) analysis (top), IHC staining of Caspase-3 (middle), and TUNEL analysis (bottom) of the tumors after different treatments, scale bar: 100μm. (F) Quantitation of caspase-3 positive cells apoptotic cells. (G) Quantitation of apoptotic cells from TUNEL staining. Percentages of cleaved caspase-3 positive cells and TUNEL-positive were quantitated by counting 100 cells from 5 random microscopic fields. Data are presented as the mean SEM, n=5. ** p<0.01, *** p<0.001.

### *In vivo* safety profile of Rod JBNps

Finally, we conducted a safety evaluation of Rod JBNp treatment. Histopathological analysis of other organs including the heart, kidney, liver, lung, and spleen was performed. No histological differences were observed in these organs in Rod JBNp group compared to the saline group, thereby suggesting that Rod JBNp did not induce noticeable damage to these organs (**Fig 8A**). Blood was collected for the complete blood count (CBC), which included a white blood cell (WBC), red blood cells (RBC), hemoglobulin (HGB), platelet (PLT), neutrophil (NEU), lymphocytes (LYM), monocytes (MON), and hematocrit (HCT) and analyzed relative to the pre-injection baseline. Differential CBC analysis revealed that the Rod JBNps did not induce changes in all levels while Doxosome shows some significant changes in increase in level of NEU and PLT (**Fig 8B**). The Rod JBNp group exhibited a normal level of body weight during the treatment stage (**Fig 8C**). In addition, the serum blood biochemistry parameters of serum aminotransferase (ALT) and aspartate aminotransferase (AST) were also analyzed to assess liver toxicity. After Rod JBNp treatment, there were no significant changes in ALT or AST levels compared to saline (**Fig 8D and Fig 8E**). Additionally, we assessed serum IgM and IgG antibody levels after treatment relative to pre-injection baseline antibody levels via enzyme linked immunosorbent assay (ELISA). Although treatment with Doxosome resulted in the elevation of adaptive immune responsive cytokine IgM, the Rod JBNp treatment regimen did not increase IgM or IgG inflammation levels (**Fig F** and **Fig 8G**). Together, these results demonstrate a better safety profile of Rod JBNp when compared to the Doxosome groups.

**Fig. 8:**
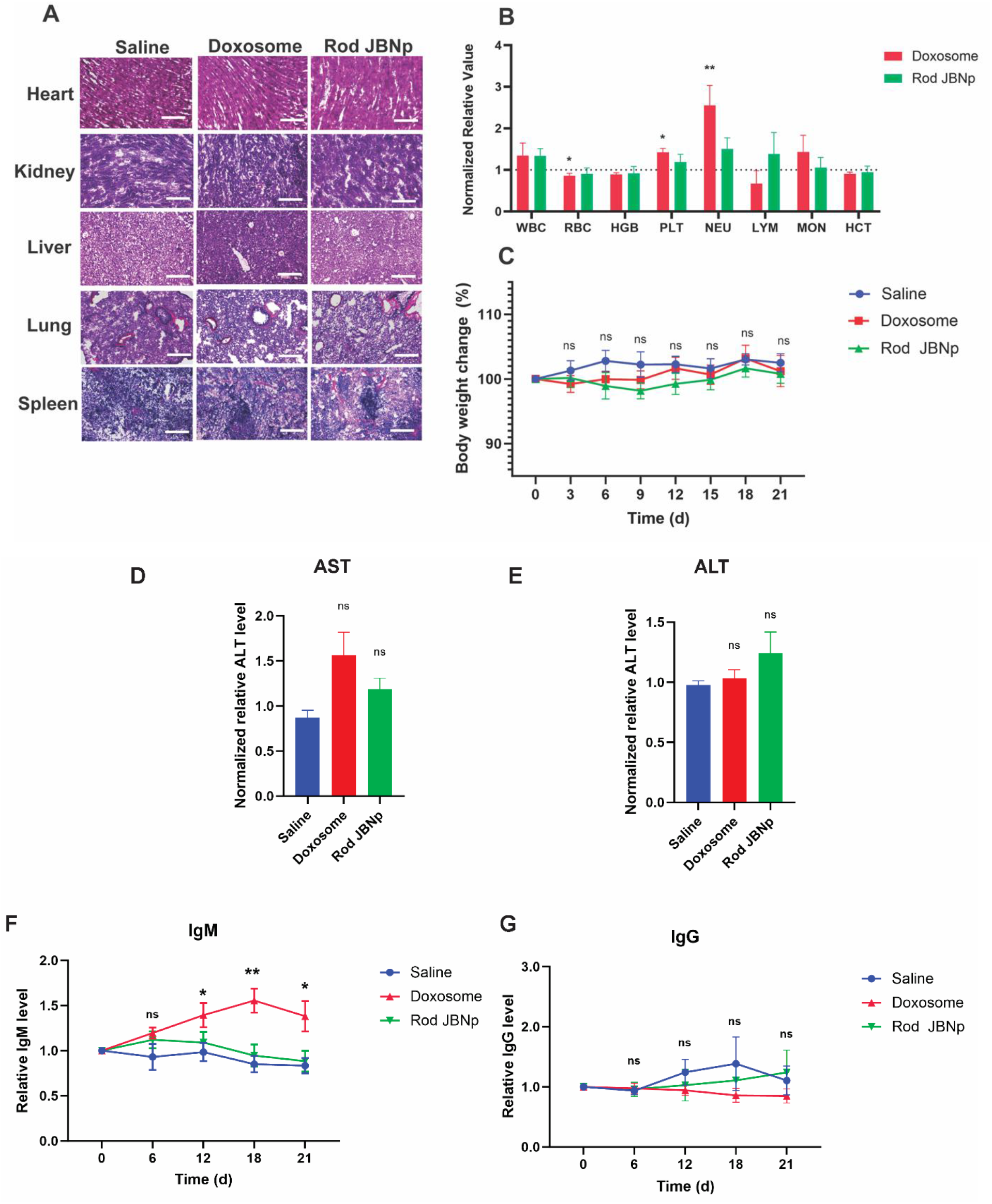
Rod shaped JBNps had a better safety profile compared to spherical Doxosome. (A) Major organs were obtained for H&E staining. Scale bar = 100 *μ*m. (B) Complete blood count (CBC) of Rod JBNP; Normalized relative value to before injection: white blood cells (WBC), red blood cells (RBC), hemoglobulin (HGB), platelet (PLT), neutrophil (NEU), lymphocytes (LYM), monocytes (MON), and hematocrit (HCT), n=5 animals. (C) Body weights change of mice in four treatment groups. (D) Level of serum aspartate aminotransferase (AST). (E) Alanine aminotransferase (ALT) analysis. (F) Serums were obtained to detect inflammatory factors IgM. (G) IgG. Data are shown as mean ± SEM (n=5 per group), *, p<0.05; **, p<0.01; ***, p<0.001.

## Discussion

In this study, we have introduced for the first time a new family of rod-shaped delivery vehicles, called Rod JBNps, inspired by JBNt and CAD methodologies. The JBNp represents a groundbreaking class of materials, distinct from existing delivery systems in several key aspects. Unlike carbon nanotubes, JBNps are characterized by their unique DNA-mimicking chemical structure [18–20]. Furthermore, while they draw inspiration from DNA, JBNps diverge from DNA origami in that they do not comprise native DNA base pairs nor possess the conventional sugar-phosphate backbone; thereby, they are resistant to enzymatic cleavage [21, 22]. In contrast to polymers, JBNps are assembled from thousands of small units through non-covalent interactions, giving them a distinct structural identity [23–26]. Lastly, while JBNps and LNPs both incorporate RNA cargos, the mechanisms of incorporation are fundamentally different. JBNps entangle RNA around their nanotubes, effectively packing the RNA within bundles to form nano-rods, whereas LNPs typically encapsulate RNA within lipid-based spherical structures.

In this research, we have employed an advanced high-throughput CAD system to model the JBNt and optimize their surface chemistry for the efficient encapsulation of cargoes. The MD simulations revealed that JBNts, assembled using G^C Janus base monomers at a vertical distance of 4 Å and a torsion angle of 30° between layers, exhibit a stable three-dimensional morphology. This specific configuration minimizes steric hindrance and maximizes stacking interactions, enhancing self-assembly. SAXS analysis confirmed an interlayer distance of 3.41Å, indicative of π-π stacking, consistent with Hicham Fenniri et al.’s previous findings of similar tubular structures. We then explore the PMF in relation to the loading of cargoes into JBNt. Our findings reveal that Lys-JBNt under high pH conditions, reduces the free energy required for cargo intercalation into Lys-JBNt. SAXS analysis corroborated these findings, confirming successful cargo loading into the JBNt. Following its validation, RSM has been effectively utilized to enhance encapsulation efficiency, eliminating the need for repetitive trials to identify the optimal method [34–37].

We then examine how Rod JBNps compare with spherical nanoparticles such as LNP, in terms of delivery capabilities in penetrating thick tissue. In contrast to traditional spherical nanoparticles like Doxosome (Aspect Ratio, AR = 1), our Rod-shaped Janus Base Nanoparticles (Rod JBNps) with an AR of 5.3 demonstrate an enhanced ability to penetrate dense ECM. [38–40]. Considering the significant role of surface charge in the endosome, the apparent pKa is another critical factor in determining the release mechanism [41, 42]. Under caveolae-mediated endocytosis (pH 6.0-7.4), the amine groups in JBNts bind to protons, exhibiting a “proton sponge” effect. This introduces positive charges to Rod JBNp, leading to the release of small molecule drugs through charge repulsion. As this was a study that focused on the development of materials and mechanism characterizations, we conducted proof-of-concept experiments. These revealed that Rod JBNps more effectively delivered DOX than Doxosome, resulting in significantly enhanced antitumor efficacy. Notably, Rod JBNp did not significantly alter CBC levels. In contrast, Doxosome administration resulted in increased neutrophil and platelet counts, a known side effect of anthracycline [43, 44]. Elevated neutrophil levels are associated with reduced cardiac function, vascular disruption, and increased collagen deposition in the heart post-DOX treatment [45]. Furthermore, our investigation into the immunogenic response revealed that repeated injections of Doxosome elevate blood serum IgM concentrations, potentially diminishing the antitumoral effect due to opsonization during circulation. [46, 47]. The excellent biocompatibility of JBNt materials seen in this study is due to the non-covalent biodegradability of the structures and DNA-mimicking chemistry. In summary, this study not only introduces the novel Rod JBNps but also comprehensively demonstrates their superior performance compared to traditional delivery systems, leveraging the innovative use of CAD for their development.

## Conclusion

In conclusion, we developed a new family of rod-shaped delivery vehicles, Rod JBNps, which surpass traditional spherical nanoparticles in penetrating such tissues. These JBNp nanorods, created from JBNts, with intercalated cargoes, were designed using a CAD methodology, integrating molecular dynamics and response surface methodology for precision and efficiency. We applied it to an ovarian cancer model as proof-of-concept and showed that Rod JBNps significantly improved ECM penetration, enhancing treatment efficacy over Doxosome. This research pioneers an advanced rod-shaped delivery vehicle for effective tissue penetration and establishes a CAD-based approach for innovative material design in drug delivery.

## Materials And Methods

### Materials

JBNt monomer was synthesized according to previously reported procedures and purified using high-performance liquid chromatography (HPLC). Doxorubicin hydrochloride (DOX·HCl) was purchased from TCI America. DOX free base was purchased from MedKoo Biosciences. Tert-butanol (*t*-BuOH) was purchased from Alfa Aesar. Methanol-d4 (CD_3_OD) was purchased from ACROS Organics. NaOD in D_2_O was purchased from Sigma Aldrich. Fibroblast, SKOV3, and MCF-7 cells were obtained from ATCC. DMEM cell culture medium (Gibco), Trypsin-EDTA (Gibco, 0.25%), Fetal Bovine Serum (FBS, Gibco), Non-essential Amino Acids Solution (100X, Gibco), Phosphate Buffered Saline (PBS, Gibco), Ethanol (70% solution) and, Lipofectamine 2000 were obtained from Thermo Fisher Scientific. Triton X-100 (Invitrogen™, 1.0%), DAPI, and 4 % formaldehyde (Invitrogen) were purchased form Fisher Scientific. Single walled carbon nanotubes dispersion (SWCNs) was purchased from US Research Nanomaterials. Polyethylenimine (PEI, branched, MW 70000, 30% w/v in water) was purchased from Alfa Aesar, CCK8 and Poly-L-lysine (PLL, 0.1% w/v in water) were purchased from Sigma. 24-well plate (Corning), 96-well plate (Corning) were purchased from Fisher Scientific, of which catalog number are 07-200-740 and 07-200-91, respectively.

### Computation-aided model setup

MD simulations were conducted for two computation-aided model systems. The first model aimed to establish the optimal condition regarding the relative positioning of monomers within the ring. This model was implemented in a simulation box of 10.0 × 10.0 × 7.2 nm^3^ consisting of water molecules having a density of 1.0 g cm^−3^. Subsequently, two G^C Janus base monomers were randomly inserted into the box under neutral conditions with their ring plane parallel to *x*-*y* plane, and the out-of-plane torsion motion was restrained. The outcome of this model was to describe the 2D-potential energy surfaces of relative stacking energies between two G^C Janus base monomers. The second model aimed to determine the energy of DOX loading to the different JBNt structures. The system included a simulation box of 10.0 × 10.0 × 7.2 nm^3^ composed of water molecules having a density of 1.0 g cm^−3^, a single DOX molecule, a JBNt structure with/without chloride ions. Four different JBNt structures were used in this model including Gly-JBNt, AA-JBNt, and neutral/acidic Lys-JBNt. The JBNts were initially made up of 16 layers along *z*-direction with a layer spacing of 4.5 Å, in which each layer was assembled by six G^C Janus base monomers to form a ring (**Fig S2A, i-ii, Fig S2B, i-ii, Fig S3A, i-ii, Fig S3B, i-ii**). To study the effect of pH values on the delivery of DOX to the JBNt, neutral form of lysine-based monomer (**Fig S3A)** and neutral DOX molecule (**Fig S4A)** were implemented in the high pH condition. Under acidic condition (low pH), the protonated lysine-based ring (**Fig S3B)** and protonated DOX form were introduced (**Fig S4B**). Additionally, chloride ions were used to neutralize the system with protonated structures before running the DOX loading simulations.

### MD simulation setup

All-atom MD and metadynamics simulations were performed using GROMACS package with CHARMM27 force field, and TIP3P explicit solvent model at room temperature (300 K) [50–53]. For each system, the steepest descent algorithm was first employed to minimize the whole system within 50,000 steps. Before the production runs, the sequential equilibrium processes in 0.5 ns-NVT (constant number of atoms, volume and temperature), 0.5ns-NPT (constant number of atoms, pressure and temperature), and 1.0 ns-NVT ensembles were performed to adjust the systems into the desired temperatures, volumes and pressure. To fully relax the JBNts before implementing DOX loading process, an additional 10.0 ns production run was performed under NPT ensemble. The leap-frog integrator was used to produce an unbiased MD trajectory for each system with an integration time-step of 1 fs under different ensemble conditions. Parrinello-Rahman barostat was employed to control the pressure at 1 bar with a coupling constant of 2 ps [54]. V-rescale was used to monitor the system temperature at 300 K with a time constant of 1 ps, which was a modified Berendsen thermostat [55]. Particle Mesh Ewald method was employed to consider the electrostatic interactions with a real-space grid distance of 1 nm [56]. The cutoff of non-bonded interactions, including electrostatic and van der Waals forces, were both set to 1.0 nm. The SETTLE algorithm was used to constrain water molecules and all non-water bonds were constrained using the LINCS algorithm [57]. For metadynamics, the out-plane torsion and in-plane translation of G^C Janus base monomer were restrained. All simulations were conducted under periodic boundary conditions. Snapshots during the simulation were rendered using the Visual Molecular Dynamics (VMD) software [58]. To model the system under low-pH environment, we adopted protonated lysine-based rings (**Fig S3B, i-ii**) and protonated forms of DOX (**Fig S4B, i-ii).** Chloride ions were introduced to neutralize the system upon the introduction of protonated Lys-JBNt and protonated DOX into the box, thus generating the required low-pH mimicking condition for the study. Following this, PMF calculations were implemented to examine the loading of DOX onto the protonated Lys-JBNt. As previous PMF calculations for Lys-JBNt were conducted under neutral conditions (pH = 7), our current focus is on evaluating the capacity of DOX loading onto the Lys-JBNt under acidic conditions.

### Metadynamics simulations

Metadynamics is an improved sampling technique used to explore the free energy profile of a system along the reaction coordinates, usually named collective variables (CVs), which can speed up the system out of a local free energy minimum and traces the other free energy minima by intermittently adding a history dependent biasing potential energy that impels the dynamics to explore those undiscovered conformations [59–62]. Compared with the motion equation of MD simulations, metadynamics has the additional forces imposing on atoms originated from the history dependent biasing potential which can be accumulated as Gaussian functions,

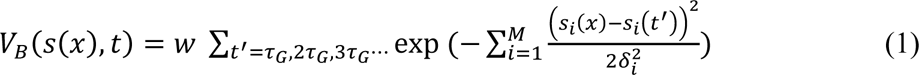

where *s*_*i*_(*x*) is the *i*^th^ of *M* reaction coordinates or collective variables. The constant *w* and *δ*_*i*_ are respectively the height and width of the Gaussian function adding at a constant time intervals of *τ*_*G*_. Well-tempered metadynamics is another improved metadynamics, which corresponds to changing the constant height of the Gaussian function as a time-dependent variable in equation 1, expressed as,

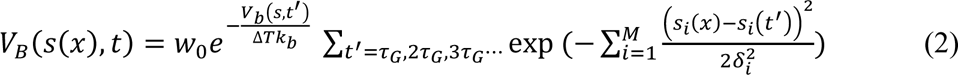

where *w*_0_ defines the initial height of Gaussian functions and Δ*Tk*_*b*_is a characteristic energy [63]. During the simulation evolution, more and more Gaussians functions are summed up and not updated until the full energy landscape is explored. Thus, over a long simulation time, the exact free energy can be evaluated from the converged biasing potential as

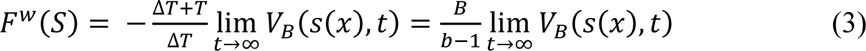

where *B* = (*T* + Δ*T*)/*T* is a scaling factor from a higher temperature *T* + Δ*T* to the normal temperature *T*.

In this work, to obtain the 2D free energy profiles, we performed the metadyamics simulations [59–61] for the first system using PLUMED 2.7 which is patched to GROMACS 2021 [64, 65]. To investigate JBNt self-assembly, two collective variables were chosen. A collective variable (CV1) was defined as the vertical distance between two G^C Janus base monomers, and another collective variable (CV2) was considered as the relative torsion angle between two G^C Janus base monomers (**Fig 2A, i**). We analyzed the variation in stacking energies as a function of the vertical distance and the relative torsion angle between two G^C Janus base monomers within a neutral condition. The Gaussian widths were set to 0.05 for CV1 and 0.05 for CV2 with the initial height of W = 0.5 kJ mol^−1^ added by every 20 ps. The bounds of the grid were set to 0 ∼ 7.0 nm for CV1 and –π ∼ +π rad for CV2. The grid bins were divided into 500 for distance space and 600 for dihedral space. To avoid overfilling, the BIASFACTOR parameter was set to 10 at TEMP = 300 K, and the Gaussian deposit time *τ*_*G*_ was set to 1 ps. A total simulation time of 20 ns was performed for metadynamics to ensure the convergence of free energy profile.

### Potential of mean force

PMF calculations were performed for the second system according to the literature using GROMACS [66]. During the delivery process of DOX, the PMF was calculated as a function of DOX−JBNt distance using umbrella sampling method [67]. The JBNt was placed in a simulation box of 10.0 × 10.0 × 7.2 nm^3^. Water molecules with the density of 1.0 g cm^−3^ were filled into the box. Subsequently, we ran a 10 ns NPT equilibrium to fully relax the JBNt (**Fig S2A, iii, Fig S2B, iii, Fig S3A, iii, Fig S3B, iii**). After that, a single DOX molecule was introduced into the simulation box, followed by the PMF simulation of DOX loading to each of JBNt structures. A harmonic potential was acted on the center of mass of DOX molecules at the *i*^th^ window along the radial direction of JBNts, and a series of separate umbrella simulations were performed. The harmonic potentials are represented by 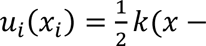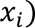, where *k* is the force constant, *x* is the coordinate of DOX along the radial direction, and *x*_*i*_ is the center position of the *i*^th^ window. In this work, the force constant was set to 1,000 kJ mol^−1^ nm^−2^, and the elongation rate of −0.3 nm ns^−1^ was chosen as the imaginary spring attached to the pull groups. The initial center position of DOX was located about 4.0 nm from the axis center of JBNt, and the final position was located about 0.5 nm from the axis center of JBNt. Window space was chosen 0.05 nm to ensure the overlapping of histogram of the positions of JBNt throughout the pulling direction (**Fig S5**) such that a continuous free energy profile could later be derived from these simulations. For each window, an NPT equilibrium run was conducted for 5 ns, followed by a production run of 5 ns for the PMF calculations. The weighted histogram analysis method (WHAM) was employed to carry out the result analysis [68]. Statistical errors were estimated with bootstrap analysis using 200 bootstraps and 50 bins for the PMF profiles [69].

### Small Angle X-ray Scattering (SAXS)

Two dimensional SAXS measurements were conducted at Brookhaven National Laboratory, utilizing the 16ID-LiX Beamline which possesses a synchrotron energy of 13.5 keV. Two aqueous solutions of JBNt (10 mg/mL) were prepared at neutral pH in the absence and presence of DOX. The samples were loaded in glass cells and maintained at 20°C during the measurements over an exposure time of 3 seconds. Radial integration was carried out to yield 1-D reduced data, the scattered intensity as a function of the scattering vector, 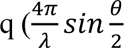, *θ* and *λ* are the scattering angle and wavelength, respectively) between 0.006 and 3.2 Å^−1^. The reduced 1-D data sets were then corrected by the background subtraction using SAXS data of water in the same cell. The corrected data were further best fitted using a combined model of core-shell cylindrical (CSC) model and a Gaussian peak in the *q* range between 0.006 and 1.3 Å^−1^.

### Response surface methodology

The following factors were analyzed at these discrete values: pH : 2.9, 6.5, 8.3; Temperature (°C) : 4°C, 25°C, 38°C; Time (hours) : 0.5, 5, 24, 48, 96. From the experimental results, a second order polynomial function was generated to model the relationship between independent variables pH (X_1_), Temperature (X_2_), and Time (X_3_) to the dependent variable, Encapsulation Efficiency (Y):

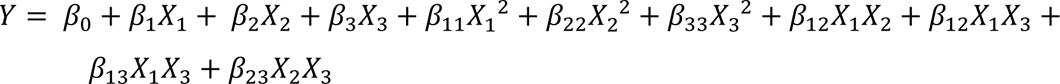

Where *β*_0_ represents the intercept, *β*_1,2,3_ represent the principal effects of each factor, *β*_11,22,33_represent the squared effects of each factor, and *β*_12,13,23_represent the interaction effect between factors. A statistical analysis, including ANOVA and multiple response prediction, was performed to optimize the model’s response. A 4D scatterplot was created to visualize the experimental data. The creation of this regression model, its corresponding statistical analysis, and the 4D experimental scatterplot were produced with Minitab ® Version 21.1 (Minitab LLC, State College, PA). An additional 4D scatterplot was created, using the RSM model instead of experimental values, in Wolfram Mathematica version 13.2 (Wolfram Research, Champaign, IL). Three 3D surface plots were generated from the RSM, each demonstrating the relationships between two factors and the response (**Fig S7**). These were created in OriginPro 2024, Package 10.1.0.170 (OriginLab, Northampton, MA).

### Fabrication of JBNt-DOX

The G^C units of JBNts were synthesized as in ref.11. The molar ratio between nitrogenous base and amino acid in JBNts is 1:1. 200 μg DOX free base was added into 1 mL JBNts solution (1 mg/mL) and vibrated with a LP Vortex Mixer (Thermo Scientific) for 24 h. And NaOH was used to adjust the pH. The solution was aged for another 1 day to allow for the drug loading process. Then the JBNt-DOX suspension was divided evenly into two parts. One part of the suspension was named as the JBNt-DOX group for the test directly. Another part of the suspension was sonicated with a sonicator (Qsonica) for 2 mins 30 secs (20 kHz, 500 W). Then it was named as the Rod JBNp group. Another 100 μg DOX free based was added into 0.5 mL PBS and vibrated with a LP Vortex Mixer (Thermo Scientific) for 24 h as well. Then the suspension was centrifuged with a Corning Mini Centrifuge for 20 seconds.

### Nuclear Magnetic Resonance experiments

JBNt monomer was first dissolved in CD_3_OD and aged for 1 day. *t*-BuOH was added as an internal standard. DOX·HCl was added in a 1:10 (DOX: JBNt monomer) mole ratio and NaOD in D_2_O was used to adjust the pH. The solution was aged for 24hr to allow for the drug loading process. Samples were measured using Bruker AVANCE 500 spectrometer at different temperatures and timepoints, respectively. Drug loading rates were calculated by peak integrations of DOX monitored at 5.5 ppm while calibrating the internal standard’s peak integration to 1 (**Table 1**).

### Characterization of Rod JBNp

Samples were measured using the Malvern Zetasizer Nano ZS90 and Nanodrop One^C^ microvolume UV-Vis spectrophotometer for DLS and UV-Vis experiments, respectively. For TEM, 3 µL of the sample was loaded on a 300-mesh copper grid and negatively stained with 50 µL 0.5% uranyl acetate and dried using filter paper. TEM images of JBNt-DOX varying sonication power and time were captured by FEI Tecnai 12 G2 Spirit BioTWIN microscope at 120 kV (**Fig S9**). The agarose gel electrophoresis assay was conducted using a 0.8% low-melting agarose gel followed by electrophoresis at 100V for 60min.

### Apparent *pK*_*a*_ assay

The apparent *pK*_*a*_of Rod JBNp was determined using a fluorescent probe 2-(p-toluidino) naphthalene-6-sulfonic acid (TNS). In brief, the TNS reagent (Sigma) was prepared as a 100*μM* stock in RNase free water. Rod JBNp were diluted in 90 *μ*L of buffered solutions (triplicates) containing 10mM HEPES, 10mM 4-morpholineethanesulfonic acid, 10mM ammonium acetate, 130 mM NaCl, where the pH ranged from 2.71 to 11.5. 10 *μ*L of stock TNS was added to the Rod JBNp solutions and mixed well in a black 96-well plate. Fluorescence intensity was monitored in a microplate reader (Molecular Devices) using excitation and emission wavelengths of 321 and 447 nm. With the resulting fluorescence values, a sigmoidal plot of fluorescence versus buffer pH was created. A sigmoidal best fit analysis was applied to the fluorescence data and *pK*_*a*_of each Rod JBNp was calculated as the pH value where half of themaximum fluorescence is reached.

### Cells and spheroid culture

Human fibroblast (ATCC) and SKOV-3 cells (graciously donated from the Dr. Hoshino lab) were cultured on T75 flasks (Fisher Scientific) at 5% CO_2_ at 37° C using McCoys 5A media (Thermo Fisher Scientific) supplemented with 1% L-glutamine and 10% FBS. To make the SKOV-3 spheroids, SKOV-3 cells were seeded on 96-well Nunclon Sphera round bottom plates (Thermo Fisher Scientific) at a density of 2 x 10^3^ cells/mL. Cells were incubated for 5 days to achieve compact and consistent spheroids before subsequent experiments.

### In vitro cytotoxicity assay

Four 96-well plates were prepared for the cytotoxicity assay of JBNts, Lipofectamine 2000 SWCNTs, PEI, and PLL. Each well of the 96-well plate received 100 μL fibroblast cell suspension (5000 cells). Put these plates into a 37 ^O^C incubator and pre-incubated fibroblast cells for 24 h. Each material was diluted with water into 10 μg/mL,5 μg/mL,1 μg/mL, and 0.2 μg/mL, respectively. After co-incubated with those materials for another 24 h, the absorption spectra of each group of solutions were measured with SpectraMax M3.

### In vitro drug delivery assay

Assembled Rod JBNp were immediately transferred to the SKOV-3 or MCF-7 cells and then incubated at 37 ° C and 5% CO_2_. The supernatant was used for the drug delivery test, which was named as the Dox free base group. All three groups of materials were diluted 10 times before using. 250 μL cell suspension (1 ×10^4^ cells) was seeded into one well of the 8-well chambered coverglass. 5 wells of cells were prepared for the test. The cells were incubated in an incubator (37 °C, 5 % CO_2_) for 24 h. Each well of cells received 250 μL of materials. For the control group, the same volume of PBS was used instead of the materials. Then the cells were co-incubated with materials for another 24 h. After incubation, DAPI was used to stain the nucleus of the cells. Nikon A1R Spectral Confocal was utilized to obtain the confocal images and Image J was used to analyze the images. Spheroids were generated from SKOV3 and cultured for approximately 5days before they reached a diameter of approximately 150 um (**Fig S15)**. Assembled Rod JBNp were immediately transferred to SKOV-3 spheroids and then incubated for 24h at 37 °C and 5% CO_2_. Then, spheroids were fixed with 4% formaldehyde, and stained with DAPI (overnight). A Nikon A1 confocal laser-scanning microscope was used for fluorescence imaging and were visualized using CLSM Z-stacks with 25um Z-intervals.

### Cell uptake mechanism study

For cell uptake mechanism studies, cells were exposed to several different concentrations of the inhibitors for 1h, pretreated with Cpz hydrochloride (100*μ*M for 30min), M*β*cd (1mM for 30 min), Cytd (4*μ*M for 1h), and Lat (2*μ*M for 30min).

### Apoptosis assay

Cells (5 × 10^5^ cells) grown on a 24-well plate for 24h were treated with DOX-free base, Non-rod JBNp, or Rod JBNp. Upon washing twice with PBS, cells were digested with 1% EDTA at 37 °C for 5min. For the cancer spheroids, the harvested spheroids following treatment were further dissociated. The cells were collected by centrifugation at 1000 rpm for 4 min. Apoptotic cells were determined with an FITC Annexin V Apoptosis Detection Kit (BD Biosciences, USA) according to the manufacturer’s protocol. Briefly, Annexin V-FITC and propidium iodide (PI) were added to the SKOV-3 cell suspension for 20 min at room temperature in the absence of light. Then, SKOV-3 cells were analyzed on a Beckman Coulter CytoFLEX flow cytometry (∼5.0 × 10^3^ cells were counted per event, **Fig S18**). For Caspase 3/7 staining, apoptotic cells were determined with an Incucyte Caspase 3/7 Dyes kit. Then, activation of apoptosis using Caspase 3/7 dye (red) and nuclei (blue) was imaged by confocal laser microscope after treatment for 48h.

### Animal studies

Mouse studies were approved by the Institutional Animal Care and Use Committee of the University of Connecticut, Health Center, where the studies were performed under Protocol A-AP-200366-9224. Female 8- to 10-weeks-old NU/J (strain : 002019) mice from the Jackson Laboratory (Bar Harbor, ME, USA) were used for all mouse studies, with random assignment of mice to experimental groups. Mice were held in a pathogen-free environment and fed at the conditions of 25 ± 2 °C and 55% humidity with a 12h light/dark cycle in accordance with guideline established by the Institutional Animal Care and Use Committee (IACUC).

### In vivo biodistribution study

Female 6- to 8-week-old NU/J mice were inoculated subcutaneously with 5 × 10^6^ SKOV-3 cells supplemented with Matrigel (Corning, #354234), and allowed to grow for 4 weeks until the tumor size was 200 mm^3^. Fluorescence imaging was performed using IVIS Spectrum Imaging system (PerkinElmer, USA) after mice were injected with 0.86 *mg*/kg of Doxosome, Non-rod JBNp, and Rod JBNp (**Fig S20**).

### In vivo Antitumor Efficacy of Rod JBNp in SKOV-3 Tumor Models

Groups with similar tumor size were assigned before the start of treatment based on initial tumor measurements using calipers. The experimenter measuring tumor size over time was blinded to group assignments. For SKOV-3, mice (n=6) were intravenously injected four times at three-day intervals with 150uL of treatment groups per mice per dose. After two months, mice were separated randomly into the following treatment groups: Saline, DOX-HCl, Doxosome, and Rod JBNp treatment (n=6). All the treatments were administered via intravenous injection every three days for 21 days. Mice were treated with Doxorubicin dose of 0.86 *m*g/kg. Tumor growth curves were plotted based on the calculation of volume = ½ (length × width^2^).

### Immunohistochemistry analysis of tissue sections

Tissue was collected and cryosections (8 *μ* m) were prepared for immunofluorescence as described previously. The rabbit anti-CD31 and anti-active caspase antibodies were used to identify endothelial and caspase-3 activity, respectively. Incubation with primary antibody was done at 4 °C overnight with either anti-CD31 diluted 1:100 or anti-active caspase diluted 1:100. After primary antibodies, tissues were incubated with the goat anti-rabbit mouse antibodies labeled with AF488. To examine the TUNEL staining, a Click-iT Plus TUNEL Assay with Alexa 647 (Molecular Probes C10619) was used. Fluorescence images were captured by Nikon A1R confocal microscope. Analysis was performed using Image J software. Cryosectioned tissues were stained with H&E for general histologic examination. And a board-certified veterinary pathologist performed blinded histopathological analysis on all sections.

### Whole blood and serum chemistry analysis

Whole blood and serum were collected from the mice via the previously indicated timeline. The peripheral blood was collected in Mini-collect K2E K2EDTA blood sample tubes. Collected samples were analyzed by IDEXX BioAnalytic for clinical chemistry and hematology of the peripheral blood. Result of the CBC studies were shown in **Table 2**.

### Statistical analysis

Statistical analyses were performed by using GraphPad Prism 7 software. Error bars were expressed as the mean ± SEM. Numerical data were analyzed via Student’s *t* test, followed by ANOVA. P values <0.05 were considered significant.

## Acknowledgements

This study is funded by NIH 7R01AR072027, NIH 1R21AR079153-01A1, NSF Career Award 1905785, NSF 2025362, NSF 2234570, NASA 80JSC022CA006, DOD W81XWH2110274 and the University of Connecticut. We would also like to thank NIH COBRE for Skeletal Health and Repair (5P30GM122732) for support and Dr. Kazunori Hoshino for providing SKOV3 cells to us. This research used 16ID-LiX beamline at the National Synchrotron Light Source II, a U.S. Department of Energy (DOE) Office of Science User Facility operated for the DOE Office of Science by Brookhaven National Laboratory under Contract No. DE-SC0012704 through a beamtime proposal (BAG-310949). The LiX beamline is part of the Center for BioMolecular Structure (CBMS), which is primarily supported by the National Institutes of Health, National Institute of General Medical Sciences (NIGMS) through a P30 Grant (P30GM133893), and by the DOE Office of Biological and Environmental Research (KP1605010). The schematic figure was created with Biorender.com.

## Notes

### Competing Interest Statement

Dr. Yupeng Chen is a co-founder of Eascra Biotech.

